# An evolutionary perspective of DNA methylation patterns in skeletal tissues using a nonhuman primate model of osteoarthritis

**DOI:** 10.1101/2020.07.31.231522

**Authors:** Genevieve Housman, Ellen E. Quillen, Anne C. Stone

## Abstract

**Objective:** Epigenetic factors, such as DNA methylation, play an influential role in the development of the degenerative joint disease osteoarthritis (OA). These molecular mechanisms have been heavily studied in humans, and although OA affects several other animals in addition to humans, few efforts have taken an evolutionary perspective. This study explores the evolution of OA epigenetics by assessing the relationship between DNA methylation variation and knee OA development in baboons (*Papio spp*.) and by comparing these findings to human OA epigenetic associations.

**Methods:** Genome-wide DNA methylation patterns were identified in bone and cartilage of the right distal femora from 56 pedigreed, adult baboons (28 with and 28 without knee OA) using the Illumina Infinium MethylationEPIC BeadChip.

**Results:** Several significantly differentially methylated positions (DMPs) and regions (DMRs) were found between tissue types. Substantial OA-related differential methylation was also identified in cartilage, but not in bone, suggesting that cartilage epigenetics may be more influential in OA than bone epigenetics. Additionally, some genes containing OA-related DMPs overlap with and display methylation patterns similar to those previously identified in human OA, revealing a mixture of evolutionarily conserved and divergent OA-related methylation patterns in primates.

**Conclusions:** Overall, these findings reinforce current etiological perspectives of OA and enhance our evolutionary understanding of epigenetic mechanisms associated with OA. This work further establishes baboons as a valuable nonhuman primate model of OA, and continued investigations in baboons will help to disentangle the molecular mechanisms contributing to OA and their evolutionary histories.

## Introduction

Osteoarthritis (OA) is a chronic, degenerative joint disease. It is characterized by a progressive degradation of cartilage and underlying subchondral bone within a joint (1) which leads to significant pain and functional limitations. According to the WHO, OA is present in 9.6% of men and 18.0% of women ages 60 or older world-wide. Of those affected, 80% have movement limitations, and 25% are unable to perform major activities of daily living (2). The CDC further notes that OA of the knee joint is especially prevalent (3) and is one of the leading causes of disability across the globe (4). The burden of OA on society demands that researchers identify the factors contributing to and aiding in the development and progression of this disease.

Although significant work has been done in this area, the complete etiology of OA is still unclear. This is because OA pathogenesis appears to be multifactorial, with both genetic and environmental influences (5,6). Additionally, epigenetic factors, such as DNA methylation which regulates gene expression, are now thought to play a more influential role in OA development (7,8). The investigation of human OA epigenetics in bone and cartilage has revealed thousands of differentially methylated candidate genes, but whether this epigenetic variation truly contributes to the development of OA and by which pathways remains unknown. Accomplishing such experimental research in humans is unethical. Thus, finding a suitable model organism in which tissue collection and direct OA progression assessment are possible is necessary for discovering the mechanisms involved in OA pathogenesis.

Several animal models of OA are currently studied (9,10). However, because the majority of these animals do not naturally develop OA, their abilities to inform our understanding of human OA are limited. Most animal models require transgenics, surgical procedures, drug injections, or non-invasive damage to a joint to induce OA. Even then, the physical manifestation of OA in these models only replicates certain stages of human OA (9,11). Additionally, in animal models that do naturally develop OA, such as guinea pigs, the occurrence of OA across individuals differs from that in humans. Specifically in guinea pigs, males have more consistent pathological alterations than females (11), while in humans, females have a higher occurrence of OA than males (4).

Conversely, among nonhuman primates, baboons develop knee OA naturally and at rates similar to those observed in humans. Like humans, the prevalence of severe OA in baboons is higher in females than in males (12,13). Additionally, in both species, the occurrence of OA is not an inevitable consequence of aging. For instance, at the Southwest National Primate Research Center, at least one-third of older baboons show no distal femur articular cartilage degradation (14). This is comparable to the almost one-third of human tissue donors (70-90 years old) that show no manifestations of knee OA (15).

Because nonhuman primates are phylogenetically close to humans, they can serve as important comparative disease models for humans (16). Baboons are large-bodied primates that develop and present OA in a manner similar to that observed in humans, making them a potentially more suitable model of OA than models currently used. Furthermore, in captive colonies of baboons, environmental factors can be regulated and controlled, thus enabling more detailed investigations of the molecular mechanisms contributing to OA pathogenesis than can be achieved in humans (12,14). Lastly, because of their evolutionary proximity to humans, using baboons as an animal model of OA will advance the evolutionary understanding of this disease (17), a perspective that has not been readily explored. The similar manifestations of OA in humans and baboons as compared to less similar manifestations of OA between humans and more phylogenetically distant animals (11), suggests that OA susceptibility and pathogenesis has undergone different waves of evolutionary conservation and divergence across species. Similarly, molecular processes innate to OA development and progression have likely also experienced selective pressures across evolution. Investigating how the molecular mechanisms contributing to OA in baboons compare to those known in humans should provide greater insight into the etiology and the evolution of OA.

In this study, we explore the evolution of OA epigenetics by identifying DNA methylation patterns in femoral bone and cartilage of 56 pedigreed, adult baboons, 28 with and 28 without knee OA, and by assessing whether DNA methylation variation is associated with OA in baboons and in a manner similar to that observed in humans.

## Materials and Methods

### Ethics Statement

Baboon tissue samples were opportunistically collected at routine necropsy of these animals. No animals were sacrificed for this study, and no living animals were used in this study.

### Samples

Samples come from captive colonies of baboons (*Papio spp*.) at the Southwest National Primate Research Center. These samples are ideal because many environmental factors that influence skeletal development and maintenance are controlled and consistent across individuals and because they have a tracked pedigree. Femora were opportunistically collected and stored at the Texas Biomedical Research Institute as previously described (18). Samples include sex- and age-matched skeletally healthy adult baboons (n=28) and adult baboons with severe knee osteoarthritis (OA, n=28) (Figure 1, Table S1). In baboons, as in humans, many skeletal features, such as bone shape and susceptibility to skeletal diseases, are sex and age dependent (13).

**Figure 1.**
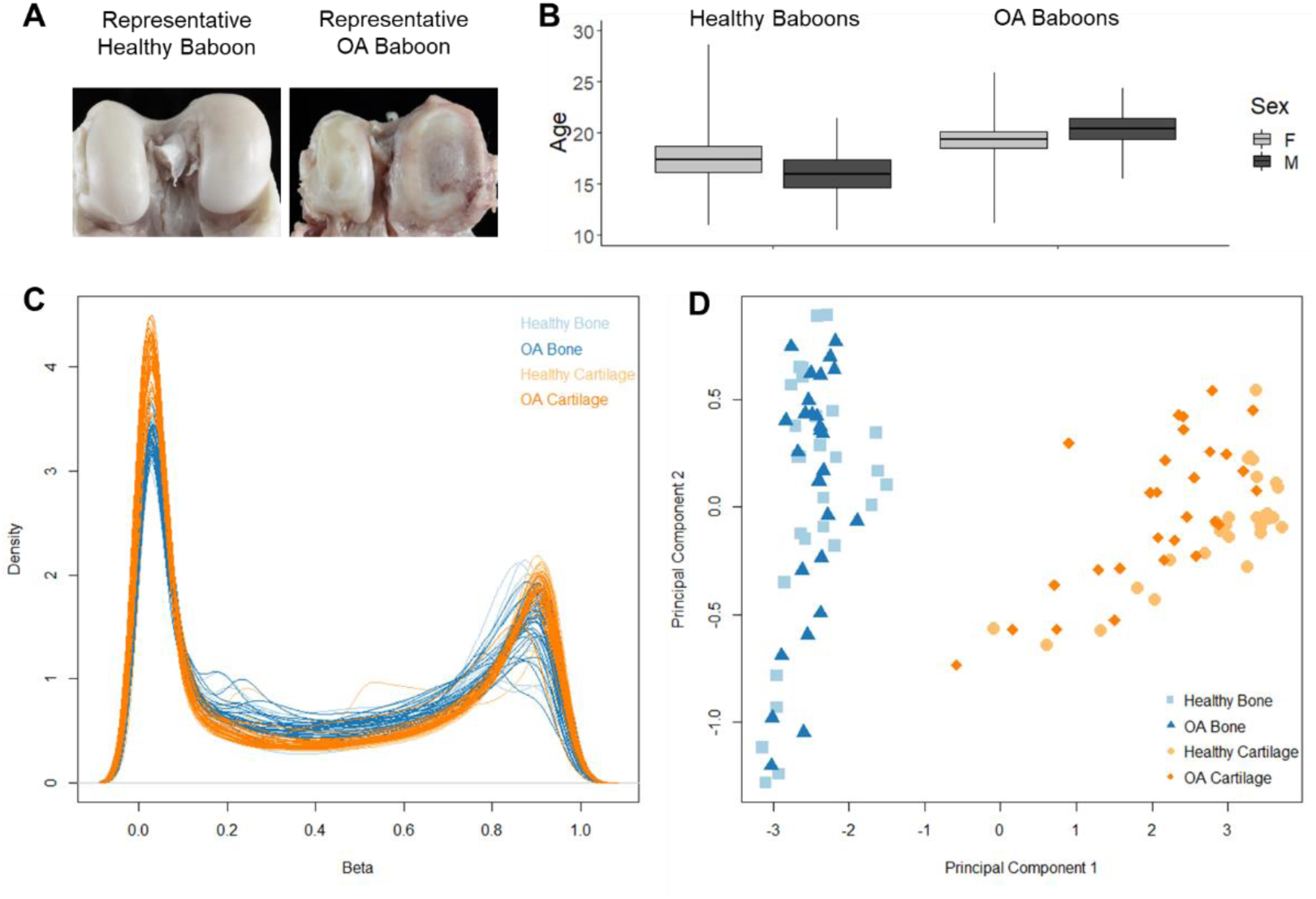
Baboon sample set and methylation data. **A**, Representative examples of baboon knees that are healthy or have severe OA. **B**, Box plots depict the average ages plus or minus one standard deviation (box), as well as the total range of ages (whiskers), for male (M) and female (F) baboons that are skeletally healthy or have OA. For males and females combined, healthy adult baboons (n=28) are 16.90±5.02 years, and OA adult baboons (n=28) are 19.73±3.41 years. **C**, Distributions of β values for each sample after normalization and probe filtering. **D**, Multidimensional scaling plot showing the first two principle components that describe methylation variation at the top 1000 most variable sites after normalization and probe filtering. Each point represents one sample that is either from healthy bone, OA bone, healthy cartilage, or OA cartilage.

### Assessment of Osteoarthritis

Baboons were classified as having healthy or OA knees as previously described (17). Briefly, each specimen was assigned an OA severity score based on macroscopic inspection of the articular surface cartilage of the distal femora: Grade 1 is unaffected, Grade 2 is mild OA (cartilage fibrillation), Grade 3 is moderate OA (cartilage lesions), and Grade 4 is advanced OA (eburnation) (12). Following this, baboons were categorized as either being healthy (100% Grade 1 on right distal femora) or having severe knee OA (variable percentage of Grades 3 or 4 on right distal femora).

### Genome-Wide DNA Methylation Profiling

For each sample, a cartilage scraping was obtained from the inferior aspect of the medial condyle on the right distal femur, and a trabecular bone core was obtained from a transverse plane through the center of the same condyle such that the articular surface remained preserved. Cartilage was collected using scalpels and processed with a homogenizer, and bone cores were collected and processed as previously described (18). Following DNA extraction, genome-wide DNA methylation was assessed using Illumina Infinium MethylationEPIC BeadChip microarrays (EPIC array) (Supplemental Text). These array data are accessible at NCBI’s GEO Series GSE103286.

### Methylation Data Processing

Raw EPIC array data were normalized and converted to β values (average methylation levels at each site) and M values (log transformed ratio of methylated signal to unmethylated signal) using standard methods (Supplemental Text). Site-specific methylation data were excluded from downstream analyses if the probe targeting the site failed detection, was classified as cross-reactive, contained SNPs at the CpG site, detected SNP information, detected methylation at non-CpG sites, targeted a site within the sex chromosomes, or had sequence mismatches with the baboon genome (18–21). Baboon SNPs that overlap with targeted probes were also identified (Supplemental Text). The finalized dataset containing 191,954 probes was used in further statistical analyses (Figure 1).

### Statistical Analysis of Differential Methylation

Genomic sites with significant differential methylation between comparative groups were identified using general linear models (GLMs) that related variables of interest to the DNA methylation patterns at each site, while accounting for the effects of other biological variables, batch effects, and latent variables. Specifically, GLMs were used to estimate differences in methylation levels for each of the following contrasts: (a) healthy bone vs. healthy cartilage, (b) OA bone vs. OA cartilage, (c) healthy bone vs. OA bone, (d) healthy cartilage vs. OA cartilage, and (e) all four combinations of tissue type and disease state. Additional variables included in these GLMs were sex, age, steady state weight, known batch effects, and latent variables to mitigate any unknown batch and cell heterogeneity effects (Supplemental Text). GLM design matrices were fit to methylation data (M values) using generalized least squares. Because each baboon contributed both a bone sample and a cartilage sample, an inter-subject correlation was performed to account for these repeated measures. Lastly, for each estimated coefficient, an empirical Bayes approach was applied to test for differential methylation given a particular contrast (Supplemental Text). Significant differentially methylated positions (DMPs) for the effect of tissue type, disease state, or both were defined as having log fold changes in methylation (M values) corresponding to an adjusted p-value less than 0.05.

In order to determine whether underlying genetic differences influenced tissue- or disease-related DMPs, generalized linear mixed models incorporating kinship were also fit to methylation data (Supplemental Text). DMPs that remained significant after adjusting for kinship were further filtered to only those with at least a 10% change in mean methylation (Δβ≥0.1) between comparative groups, as these are expected to have greater biological impact than others (22). Gene ontology (GO) and KEGG pathway enrichments for DMPs were also determined to functionally characterize DMPs (Supplemental Text), and differentially methylated regions (DMRs) were identified between each comparative group to confirm general DMP findings (Supplemental Text). Lastly, baboon OA-related methylation changes identified in this study were compared to OA-related methylation patterns previously identified in humans (Supplemental Text).

## Results

### Differential Methylation across Tissue Types and Disease States

Significant DMPs were interrogated from 191,954 sites and identified between tissue types (bone vs. cartilage) and disease states (OA vs. healthy), as well as among these variables in combination (Figure 2A, Table S2, FileS1). Filtering DMPs using different thresholds did not substantially change the general trends observed (Supplemental Text, Table S3, Table S4). Overall, more DMPs and DMRs were found between tissue types than between disease states, and cartilage samples revealed more DMPs and DMRs between disease states than did bone samples (Table S5, File S2). Fittingly, the variation in methylation patterns across the final set of DMPs (which remained statistically significant after accounting for kinship and had a Δβ≥0.1) clusters bone and cartilage tissue types into distinct and separate groups but does not cluster OA and healthy individuals as effectively (Figure 2B, Figure S1). While OA and healthy samples within cartilage differentiate relatively well, OA and healthy samples within bone are not clearly differentiated. This final set of DMPs which were used in downstream functional analyses included: 47,386 DMPs that differentiated healthy bone and cartilage, 48,562 that differentiated OA bone and OA cartilage, 39 that differentiated healthy bone from OA bone, 4,298 that differentiated healthy cartilage from OA cartilage, and 2,678 that were differentiated among all four combinations of tissue type and disease state.

**Figure 2.**
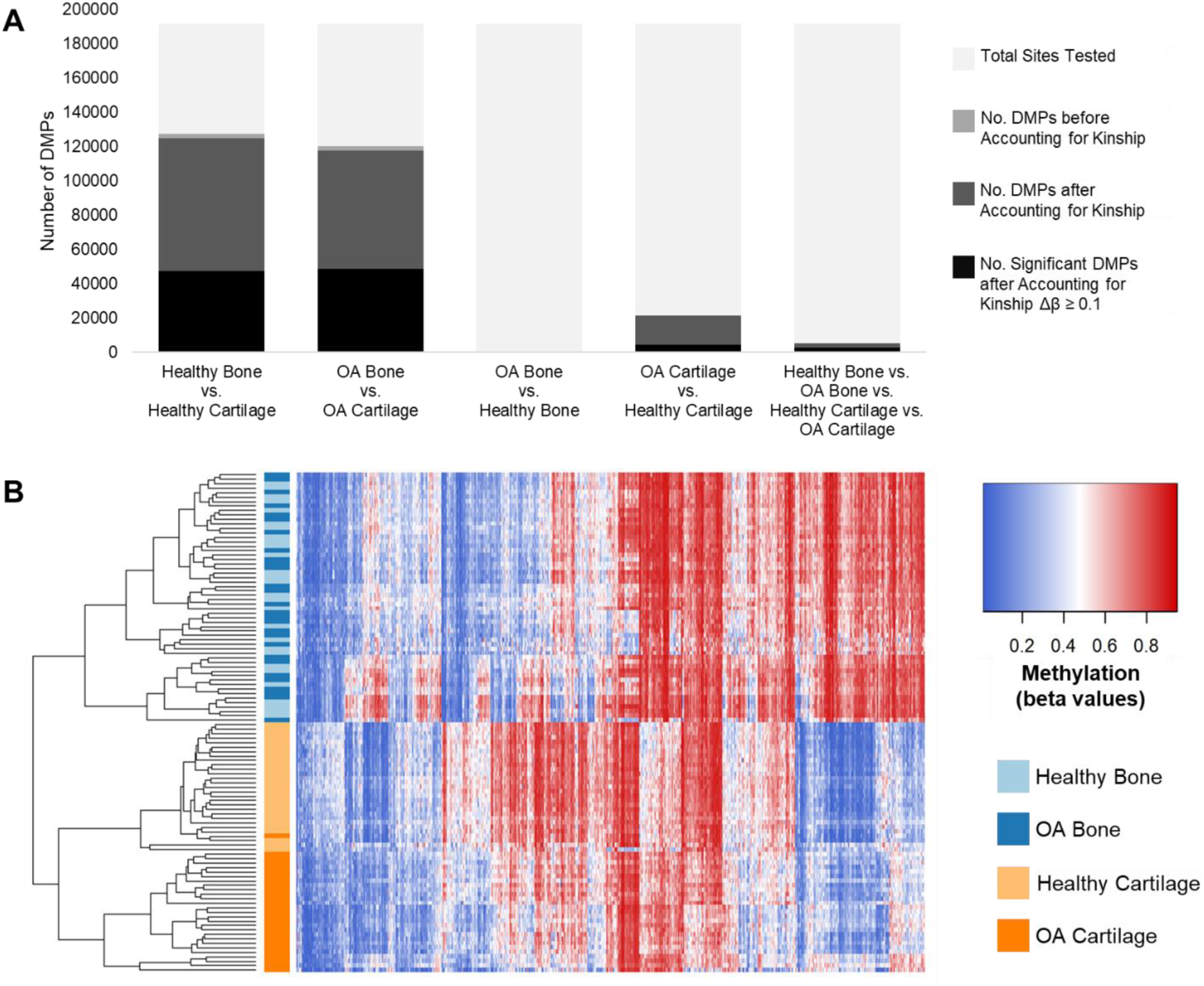
Significant DMPs. **A**, Bar chart showing the number of DMPs between comparative groups. Results include the number of DMPs that were statistically significant before accounting for kinship, the number of DMPs that remained significant after accounting for kinship, and the number of DMPs that remained significant after accounting for kinship and had at least a 10% change in mean methylation (Δβ≥0.1) between comparative groups. **B**, Heatmap depicting the DNA methylation levels (β values) of all DMPs after accounting for kinship with Δβ≥0.1 between all four combinations of disease state and tissue type (x-axis) in all baboon samples (y-axis, n=112). Red indicates higher methylation at a DMP, and blue indicates lower methylation at a DMP. The dendrogram of all samples (y-axis) clusters samples based on the similarity of their methylation patterns.

These DMPs are associated with several genes that have distinct GO biological processes (Figure S2, FileS3) and KEGG pathway functions (FileS4). Specifically, tissue-related DMPs are enriched for biological processes related to anatomical structure development and morphogenesis, biological adhesion, cell communication, and signaling, which are fitting given the roles of bone and cartilage in skeletal development and maintenance. Not enough bone OA-related DMPs were identified to detect significant functional enrichment, so only related but non-significant biological processes could be evaluated. In contrast, cartilage OA-related DMPs are enriched for several biological processes, including connective tissue development, skeletal system development, cartilage development, and ossification.

When comparing the DMPs, DMRs, GO functions, and KEGG pathways that were identified between each comparative group, distinct patterns were identified. Out of the 54,302 unique tissue-related DMPs identified across healthy and OA individuals, 77% overlap. Similarly, out of the 24,079 unique tissue-related DMRs, 71% overlap. This pattern holds for GO functions in which 73% overlap but changes slightly for KEGG pathways in which 60% overlap (Figure S3). Overall, these data indicate that the locations and functional associations of differential methylation between bone and cartilage tissues are very similar in both healthy and OA individuals. Conversely, out of the 4,332 unique disease-related DMPs identified across bone and cartilage tissues, only 0.1% are identical. Similarly, out of the 3,547 unique disease-related DMRs, 0.1% are identical. This pattern holds for GO functions in which 0% overlap but changes slightly for KEGG pathways in which 12% overlap (Figure S4). Overall, these data indicate that the locations and functional associations of differential methylation between healthy and OA individuals are different in bone and cartilage tissues.

### Overlap of OA-Related Epigenetic Patterns among Baboons and Humans

Baboon OA-related methylation changes were compared to OA-related methylation patterns previously identified in humans. While several genes and CpG sites show changes in methylation that are unique to each species (75-90% of genes and 91-99% of CpG sites depending on the joints and tissues included in the comparison) (Table S6, Table S7, File S5), loci with differential methylation unique to humans have functions that generally overlap with those identified in baboons. For example, when comparing joint- and tissue-matched samples across species, 65% of genes with human-specific differential methylation are involved in the same GO biological processes that are enriched in baboon cartilage OA-related DMPs (FileS3).

A subset of loci showing significant changes in both species overlap (10-25% of genes and 1-9% of CpG sites depending on the joints and tissues included in the comparison) (Table S6, Table S7, File S5). Of these overlapping loci, approximately half display conserved OA-related methylation patterns between species and half display divergent OA-related methylation patterns between species. For example, when comparing joint- and tissue-matched samples across species, 596 differentially methylated genes overlap, of which 237 genes have OA-related methylation changes in the same direction, 221 have reversed signals, and 138 were excluded due to conflicting methylation patterns present across different CpG sites within a gene or across different human OA studies. Genes with generally conserved patterns of OA methylation may play particularly important roles in OA pathogenesis. Thus, cases of genes that display similar OA-related methylation changes, as well as a range of reversed OA-related methylation changes, were evaluated more closely.

For example, in baboon cartilage, out of the 24 CpG sites examined in the *TBX4* gene, 8 OA-related DMPs were identified, all of which were hyper-methylated (Figure 3A, Table S8). Additionally, 2 DMRs showing OA hyper-methylation were found nearby. Similarly, OA-related hyper-methylation of *TBX4* in cartilage was observed in all human studies. Conversely, while OA hyper-methylation of *TBX4* has also been observed in human bone, no OA-related DMPs or DMRs were observed in baboon bone.

**Figure 3.**
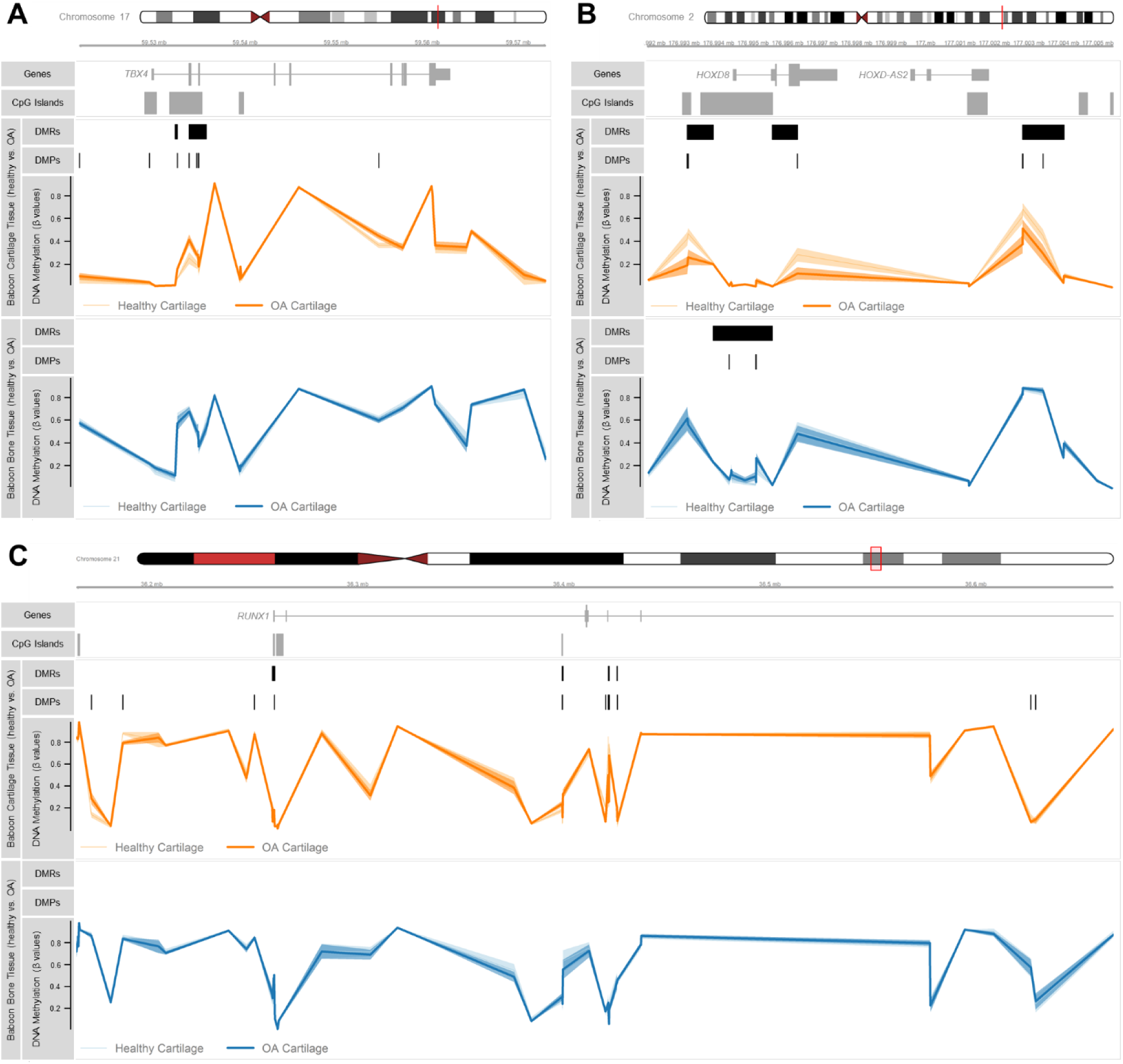
Methylation across OA-related genes. Plots show the genomic coordinates (hg19) of candidate OA genes and nearby annotated CpG islands, along with baboon cartilage and bone methylation levels (average β values and 95% confidence intervals) for all sites examined with the positions of significant OA-related DMRs and DMPs identified in bone and cartilage denoted. All DMPs displayed remained statistically significant after accounting for kinship. **A**, Plot of baboon methylation across *TBX4* (gene: chr17:59529134-59562471). *TBX4* in baboon cartilage displays similar OA hyper-methylation patterns as those observed in humans. **B**, Plot of baboon methylation across *HOXD8* (gene: chr2:176994422-176997423). *HOXD8* in baboon bone displays similar OA hyper-methylation patterns as those observed in humans. **C**, Plot of baboon methylation across *RUNX1* (gene: chr21:36160098-36421595). *RUNX1* in baboon cartilage displays a mixture of OA hypo- and hyper-methylation patterns which overlap with the OA-related hypo- and hyper-methylation observed in humans to reveal a combination of conserved and divergent methylation patterns. See Table S8, FileS1, and FileS2 for additional information.

Next, in baboon bone, out of the 26 CpG sites examined in the *HOXD8* gene, 4 OA-related DMPs were identified, all of which were hyper-methylated (Figure 3B, Table S8). Additionally, 1 DMR showing OA hyper-methylation was found nearby. Similarly, OA-related hyper-methylation of *HOXD8* in bone was observed in all human studies. While significant OA-related changes in *HOXD8* methylation have not been observed in human cartilage, several were identified in baboon cartilage and showed a reversed signal to that in bone. Specifically, in baboon cartilage, 6 OA-related DMPs were identified, all of which were hypo-methylated. Additionally, 3 DMRs showing OA hypo-methylation were identified nearby.

Lastly, in baboon cartilage, out of the 60 CpG sites associated with *RUNX1* that were examined, 16 DMPs were identified of which 9 were hypo-methylated and 7 were hyper-methylated (Figure 3C, Table S8). Additionally, 1 DMR showing OA hypo-methylation and 3 DMRs showing OA hyper-methylation were identified nearby. Similarly, a mixture of OA-related hypo- and hyper-methylation of *HOXD8* was observed across human studies. While this makes gene-level species comparisons difficult, site-specific comparisons reveal that human OA-related patterns in knee cartilage are similar to those observed in baboon knee cartilage, while human OA-related patterns in hip cartilage show reversed signals.

## Discussion

Here, we examined DNA methylation variation in femoral bone and cartilage from a baboon model of OA to assess the evolutionary conservation of epigenetic-OA associations in the primate lineage. Overall, more DMPs were found between tissue types than between disease states, and more OA-related DMPs were found in cartilage than in bone. Additionally, our study identified substantially more DMPs than a previous investigation of baboon OA (17). Although this prior study used the 450K array, the total number of sites examined was similar, suggesting that increased power was primarily due to an increased sample size. The expansion of DMP findings allowed for subsequent analyses of DMRs which showed similar number trends and gene associations, further confirming the DMP results and their functional implications.

Tissue-related DMPs clearly distinguish bone from cartilage and are highly similar across healthy and OA baboons, indicating that tissue types maintain distinct epigenetic profiles regardless of disease progression. Maintaining distinct epigenetic profiles is expected in healthy tissues, as bone and cartilage have distinct roles within the skeletal system. In contrast, OA bone and cartilage can, at a macroscopic level, look like a single, blended, pathological tissue type. In such a scenario, the epigenetic profiles of OA bone and OA cartilage are expected to be more similar than those of healthy bone and healthy cartilage. However, we instead find similar degrees and types of methylation differences across tissues regardless of disease state, and this suggests that during baboon OA pathogenesis, bone and cartilage tissues are not de-differentiating and becoming more similar to one another.

With respect to OA-related DMPs, these epigenetic patterns readily distinguish OA and healthy states in cartilage, but not in bone. This larger number of OA-related DMPs identified in cartilage than in bone may have etiological implication. For instance, it suggests that cartilage epigenetics may have a more influential role in the development of OA than bone epigenetics. This finding is reinforced by several human OA studies (23–29). Of note, this study only examined trabecular bone, so it cannot speculate on the degree to which subchondral cortical bone epigenetics may impact OA pathogenesis. Alternatively, the discrepancy between bone and cartilage may be an artifact of bone tissue being more heterogeneous than cartilage tissue, and thus, less likely to detect OA epigenetic signals using the methods performed in our study. Few OA-related DMPs or associated functions overlap between bone and cartilage, as well. In bone, functional enrichment analyses were limited by the small number of DMPs identified, but some non-significant but related biological processes include fibroblast apoptotic function and actin filament-based movement. Conversely, several gene functions are enriched in cartilage OA-related DMPs, the predominant being connective tissue development, skeletal system development, cartilage development, and ossification. Altogether, these finding reveal stronger and more functionally relevant OA-related epigenetic changes in cartilage as compared to bone. However, it is unclear whether these epigenetic changes play a role in initiating OA pathogenesis, are a byproduct of OA progression, or a combination of both.

Nevertheless, these disease-related DMPs provide further insight into the evolution of OA epigenetics. When our baboon OA findings are compared to differentially methylated loci in human OA studies, hundreds of CpG sites and genes overlap between species. Although the majority of OA-related epigenetic patterns show no evidence of species overlap, of those that are present in both baboons and humans, several display similar patterns of OA-related methylation changes across species. Because these molecular changes are shared across species when they develop the same disorder, these evolutionarily conserved OA-related epigenetic patterns may be particularly relevant for understanding molecular contributions to OA initiation and progression. Specific examples of genes with OA methylation patterns shared across species are as follows, each of which varies in its levels of conservation and potential regulatory impact.

*TBX4* displays similar OA hyper-methylation in human cartilage (23,25,26) and baboon cartilage but not in bone. *TBX4*, also known as T-box-4, codes for a transcription factor that regulates lower limb developmental processes (30). Mutations of this gene in humans have resulted in pathologies of femoral joints (31,32), areas that also readily develop OA (4,33). Thus, less severe alterations of *TBX4* via epigenetic changes are likely mechanistically involved in the chronic pathogenesis of OA, as evidenced by the conserved hyper-methylation of this gene in both human and baboon OA cartilage. Further, although the methylation changes observed in baboon cartilage are modest, they primarily occur in an upstream CpG island of *TBX4*, suggesting that they may regulate this gene’s expression.

Conversely, *HOXD8* displays similar OA hyper-methylation in humans (26,34) and baboon bone but a reversed signal in baboon cartilage. *HOXD8*, also known as homeobox D8, codes for a transcription factor that regulates lower limb morphogenesis, and lower limb malformations result from mutations in this gene (35,36). Similar to *TBX4*, less severe alterations of *HOXD8* via epigenetic changes may be mechanistically involved in OA pathogenesis. However, the evolutionarily conserved impact of methylation changes in *HOXD8* may be restricted to bone. Although substantial *HOXD8* OA-related hypo-methylation is observed in baboon cartilage, this pattern is not conserved as this gene is not differentially methylated in human cartilage. It might be the case that this epigenetic pattern has simply not yet been identified in human cartilage, possibly because human studies often compare degraded and non-degraded cartilage within the same OA joint (26,27) as opposed to cartilage from separate OA and healthy joints or because human studies often evaluate healthy cartilage of the hip joint (24,37) as opposed to healthy cartilage of the knee joint. Regardless, the moderate *HOXD8* methylation changes observed in baboon cartilage suggest that they may regulate this gene’s expression and have an important molecular function during OA pathogenesis despite not being conserved in humans. Alternatively, the small methylation changes observed in baboon bone imply a limited effect on gene regulation irrespective of their evolutionary conservation.

Lastly, *RUNX1* displays a mixture of OA-related hypo- and hyper-methylation in baboons, of which some patterns are conserved with humans and others are divergent (23,25–27,37). *RUNX1*, also known as runt related transcription factor 1, is involved in the regulation of skeletal cell development and differentiation (38). In addition to its association with OA, *RUNX1* polymorphisms show variable effects on rheumatoid arthritis (45,46,47,48,49). Thus, epigenetic changes to *RUNX1* may impact OA pathogenesis. Fittingly, previous research identified OA-related hypo-methylation of *RUNX1* in baboon cartilage (17). This same pattern was identified in our current study, along with additional regions of OA-related hyper-methylation. This mixture of OA-related epigenetic patterns across *RUNX1* persists in humans, but it is unclear whether site-specific similarities and differences across species are truly conserved and divergent. Baboon OA-related patterns do appear to be more similar to those observed in human knee cartilage than those observed in human hip cartilage which often show reversed signals. Nevertheless, the small methylation changes observed in baboon cartilage, which imply a limited effect on gene regulation, further dampen the etiological implications of *RUNX1* OA-epigenetic findings.

In addition to the subset of OA-related methylation patterns that are evolutionarily conserved between humans and baboons, we found several other associations that are not conserved. Nevertheless, loci with human-specific differential methylation have functions that generally overlap with those functions identified in baboon DMPs. Species differences in OA epigenetics may be due to general speciation events that took place during the evolution of each taxonomic group, to slight differences in the development or manifestation of OA in each species, or to artifacts of our experimental design or that of human OA studies. For instance, our study only evaluated one population of baboons; whereas, several studies of OA in different populations were considered for humans (23–29,34,44–48). This likely introduced additional noise into the human OA dataset which was not present in our baboon dataset. In order to account for this, replicate studies of OA epigenetics should be done in in different baboon populations. The sample size of our study was comparable to that of human OA studies (minimum: n=12, maximum: n=117) (23–27,34,37,44), so this likely did not impact species differences. Nevertheless, to identify candidate epigenetic alterations that underlie variation in knee OA more accurately, even larger sample sets of both baboons and humans should be considered that perhaps focus only on cartilage epigenetics. Lastly, our study used a stringent DMP cutoff threshold to limit results to biologically relevant methylation changes (22). However, while some human OA studies enforce comparable thresholds (23,24,26,37), some do not (28,34,47).

Therefore, many genes previously classified as being differentially methylated in human OA may be false positives. Until further work to identify the mechanisms through which OA naturally develops and progresses is done in humans or other model systems, the validity of currently known candidate genes with OA-related methylation will remain unknown.

In conclusion, this study further informs our understanding of DNA methylation variation in two skeletal tissues from baboons, as well as the degree to which OA affects this variation. Our results help to clarify the etiology of OA and, from an evolutionary perspective, the conservation of epigenetic mechanisms associated with OA. Additionally, this work further establishes baboons as a valuable nonhuman primate model of OA. Our findings warrant further investigation in a larger and more phylogenetically diverse sample set of nonhuman primates, as such future research will provide further insight into the evolution of OA pathogenesis.

## Author Contributions

All authors were involved in drafting or critical revision of this article, and all authors approved the final version to be published. G.H. had full access to the study data and takes responsibility for the integrity of these data and the accuracy of their analysis.

Study conception and design: G.H., E.E.Q., A.C.S. Acquisition of data: G.H.

Analysis and interpretation of data: G.H.

## Acknowledgements

This work was supported by the National Institutes of Health (P01HL028972 to Anthony G. Comuzzie); the Leakey Foundation (Research Grant for Doctoral Students to G.H.); the Wenner-Gren Foundation (Gr. 9310 to G.H.); the Nacey Maggioncalda Foundation (James F. Nacey Fellowship to G.H.); the International Primatological Society (to G.H.); Sigma Xi (Grant-in-Aid of Research to G.H.); the ASU Center for Evolution and Medicine (Venture Fund to G.H.); and the ASU Graduate Research and Support Program (to G.H.). Additionally, this investigation used resources that were supported by the Southwest National Primate Research Center grant P51OD011133 from the Office of Research Infrastructure Programs, National Institutes of Health.

We thank Eric D. Johnson and members of the Department of Genetics at the Texas Biomedical Research Institute, including Anthony G. Comuzzie, Lorena M. Havill, Anne Sheldrake, Jaydee Foster, Kara Peterson, Mel Carless, and Laura Cox, for helpful discussions.

Newly reported data have been made available on NCBI’s Gene Expression Omnibus and are accessible through the GEO Series accession number GSE103286.

