## Supplemental Text for "An evolutionary perspective of DNA methylation patterns in skeletal tissues using a nonhuman primate model of osteoarthritis"

### Supplemental Materials

### Supplemental Text

#### Additional Details on Genome-Wide DNA Methylation Profiling Methods

Skeletal tissues were collected from the medial condyle of the femur as this joint region is a common site of OA development in baboons and humans. Trabecular bone and articular cartilage were sampled because both tissues are clinically relevant with respect to disease progression. Additionally, human OA epigenetic studies are based on both trabecular bone and cartilage, so for comparative purposes, it was important to standardize tissue types.

Several considerations were made to minimize unwanted variation between samples and comparative groups. First, only baboons from captive colonies with similar environmental exposures were included in this study. Regulatory patterns in skeletal tissues may be impacted by environmental forces, such as diet, exposure to sunlight which influences vitamin D production, and biomechanical loading during daily activities. Thus, restricting samples to only those exposed to controlled and consistent environmental conditions was expected to minimize skeletal epigenetic variation due to these external forces. Second, only age- and sex-matched, sexually mature baboons were included in this study to control for the potential effects of growth and developmental processes on skeletal epigenetic patterns. Lastly, bone and cartilage tissues were collected from the same portions of baboon femora. Methylation patterns in cartilage are known to vary between joints and between different sites within a joint (1–5), and although similar studies in bone have not been conducted, epigenetic signatures are also expected to vary across locations. Thus, collecting tissues from the same portion of the femur further ensured that sample location was not a strong effect on skeletal epigenetic variation.

Genome-wide DNA methylation was assessed using Illumina Infinium MethylationEPIC BeadChip microarrays (EPIC array) which analyze the methylation status of over 850,000 sites throughout the genome. For each sample, 400ng of genomic DNA was bisulfite converted using the EZ DNA Methylation™ Gold Kit according to the manufacturer's instructions (Zymo Research), with modifications described in the Infinium Methylation Assay Protocol. Following manufacturer guidelines (Illumina), this processed DNA was then whole genome amplified, enzymatically fragmented, hybridized to the arrays, and imaged using the Illumina iScan system.

#### Additional Details on Methylation Data Processing

Raw fluorescent data were normalized to account for the noise inherent within and between the arrays themselves. Specifically, a normal-exponential out-of-band (Noob) background correction method with dye-bias normalization (6) was performed to adjust for background fluorescence and dye-based biases. This was followed with a between-array normalization method (functional normalization) (7) which removes unwanted variation by regressing out variability explained by the control probes present on the array as implemented in the minfi package in R (8,9) which is part of the Bioconductor project (10). This method has been found to outperform other existing approaches for studies that compare conditions with known large-scale differences (7), such as those assessed in this study.

After normalization, methylation values (β values) for each site were calculated as the ratio of methylated probe signal intensity to the sum of both methylated and unmethylated probe signal intensities (Equation 1). These β values range from 0 to 1 and represent the average methylation levels at each site across the entire population of cells from which DNA was extracted (0 = completely unmethylated sites, 1 = fully methylated sites). Because β values have high heteroscedasticity, they are not statistically valid for use in differential methylation analyses (11). Thus, M values were also calculated and used in downstream analyses instead (Equation 2).

**Equation 1:** $\beta \text{ Value} = \frac{\text{Methylated Signal}}{(\text{Methylated Signal} + \text{Unmethylated Signal})}$ **Equation 2:** $\text{M Value} = \log\left(\frac{\text{Methylated Signal}}{\text{Unmethylated Signal}}\right)$

Every probe on the EPIC array is accompanied by a detection p-value. Probe with failed detection levels (p-value > 0.05) in greater than 10% of samples were removed from downstream analyses. Samples in which more than 30% of the probes failed detection were also removed from downstream analyses.

Because probes on the EPIC array were designed to hybridize with human DNA, using baboon DNA in these assays further required that probes non-specific to the baboon genome be removed from downstream analyses, as they likely produce biased methylation measurements. Such probes containing sequence mismatches with the baboon genome were computationally filtered out using methods modified from (12,13) and previously described in (14). Briefly, blastn (15) was used to map the 866,837 EPIC array probes (50bp each) onto the *Papio anubis* genome (Assembly: Panu_2.0, Accession: GCF_000264685.2). Probes were kept only if they successfully mapped to the baboon genome, had only 1 unique BLAST hit, targeted CpG sites, had 0 mismatches in 5bp closest to and including the CpG site, and had 0-2 mismatches in 45bp not including the CpG site (14). This filtering retained 209,802 probes, which cover approximately 23,446 genes with an average coverage of 8 probes per gene and maintain a wide distribution throughout the genome (14).

Additionally, probes were removed if they were cross-reactive, contained SNPs at the CpG site, detected SNP information, detected methylation at non-CpG sites, or targeted a site within the sex chromosomes (8,9,16). This filtering produced a finalized set of 191,954 probes which were used in downstream analyses.

Lastly, the overlap of baboon SNPs (10.5281/zenodo.2583291) with EPIC array probes was investigated to ensure that significant methylation differences were not biased by such genetic variation. Specifically, for probes that successfully mapped to the baboon genome, the number of baboon SNPs that overlapped with the entire probe region and that occurred at the targeted CpG site were determined as previously described (14). Briefly, because less than 0.55% of baboon-filtered probes have a CpG-SNP overlap, such probes were not filtered before running statistical analyses. However, the small number of probes with CpG-SNP overlaps that were also identified as significantly differentially methylated are noted in corresponding supplementary materials (FileS1, Table S8).

#### Additional Details on Statistical Analysis of Differential Methylation

We designed and tested general linear models (GLMs) and generalized linear mixed models (GLMMs) which related variables of interest (tissue type and disease state) to the DNA methylation patterns for each site, while accounting for other fixed effects such as sex, age in years, steady state weight in kilograms, batch effects (array number and position), and latent variables, as well as other random effects such as kinship (17). The specific GLM and GLMM design matrices and contrasts used in this study were:

**Equation 3A**. methylation ~ tissue type and disease state contrasts + sex + age + weight + batch effects + latent variables
**Equation 3B**. methylation ~ tissue type and disease state contrasts + sex + age + weight + batch effects + latent variables + kinship
**Equation 3C**. methylation ~ sex + age + weight + batch effects + latent variables + kinship

| No. | Comparison | Explicit Contrast |
|---|---|---|
| **(a)** | between bone and cartilage in healthy baboons | healthy bone vs. healthy cartilage |
| **(b)** | between bone and cartilage in OA baboons | OA bone vs. OA cartilage |
| **(c)** | between healthy and OA baboon bone | healthy bone vs. OA bone |
| **(d)** | between healthy and OA baboon cartilage | healthy cartilage vs. OA cartilage |
| **(e)** | among all 4 combinations of tissue type and disease state | healthy bone vs. healthy cartilage vs. OA bone vs. OA cartilage |

First, latent variables were calculated using the iteratively re-weighted least squares approach in the sva package in R (18–21). The 14 latent variables estimated were included to help mitigate any unknown batch and cell heterogeneity effects on methylation variation at each site. Alternative methods to account for cell heterogeneity exist, but they are specific to whole blood (18,22), require reference epigenetic data, or are reference free methods (23) that are comparable to the sva method (24). Out of the known cell types in skeletal tissues (25), only chondrocytes and osteoblasts have reference epigenomes available on the International Human Epigenomics Consortium, and these are only for humans, not nonhuman primates. Thus, because no standard method is available to correct for the heterogeneous cell structure in nonhuman primate skeletal tissue, the described sva method was chosen.

Next, GLM design matrices (Equations 3A) were fit to processed and filtered methylation data (M values) using generalized least squares in the limma package in R (10,26,27), and the estimated coefficients and standard errors for the defined tissue type and disease state contrasts were computed. Because each baboon contributed both a bone sample and a cartilage sample, an inter-subject correlation was performed to account for these repeated measures (28) and included in each GLM. For each coefficient, an empirical Bayes approach was applied using the limma package in R (27,29–31) to compute moderated t-statistics, log-odds ratios of differential methylation, and associated p-values adjusted for multiple testing (32). Significant differentially methylated positions (DMPs) for the effect of tissue type, disease state, or both were defined as those having log fold changes in M values corresponding to an adjusted p-value of less than 0.05.

In order to account for genetic relatedness, the coefficient of relatedness (Equation 4), or the expected proportions of alleles that are identical by descent between two individuals, was computed for each sample from known pedigree data using the kinship2 package in R (33). Following this, two GLMMs were designed and tested using the lmekin function of the coxme package in R (34). The first GLMM (Equation 3B) regressed methylation data (M values) against the tissue type and disease state contrast effects while adjusting for additional variables (sex, age, batch effects, latent variables) as fixed effects and kinship (phi2) as a random effect (35). The second GLMM (Equation 3C) performed the same regression but with the tissue type and disease state contrast effects removed. The log likelihoods of each GLMM were compared using a chi-square test to determine which model better explained the variation in methylation. For this test, the degrees of freedom were calculated as the absolute difference in the Akaike's information criteria for each model (36). When the first GLMM containing the tissue type and disease state contrast effects (Equation 3B) performed significantly better than the alternative model (Equation 3C) (p-value < 0.05), this confirmed that the site remained a significant DMP for the effects of tissue type, disease state, or both when adjusting for the added effects of kinship. These DMPs were classified as significant when adjusting for kinship. Conversely, when the first GLMM containing the tissue type and disease state contrast effects (Equation 3B) did not perform better than the alternative model (Equation 3C) (p-value ≥ 0.05), this indicated that the site was not a significant DMP for the effect of tissue type, disease state, or both when adjusting for the added effects of kinship. In these instances, DMPs were classified as non-significant when adjusting for kinship and were no longer considered as sites with significant differential methylation.

**Equation 4:** phi2 = 2 x kinship coefficients

Further, DMPs that had at least a 10% change in mean methylation ($\Delta\beta \geq 0.1$) between comparative groups were identified, as these may have greater biological impact than others (12).

Accounting for kinship slightly reduces the DMP counts per comparative group, and applying an additional $\Delta\beta \geq 0.1$ threshold substantially decreases the final DMP counts. Across all cutoffs, more than half of tissue-related DMPs are hyper-methylated in bone as compared to cartilage, while for OA-related DMPs, only those identified in cartilage show a consistent trend across cutoffs, with over half displaying hypo-methylation in OA baboons as compared to healthy baboons (Table S3). Additionally, the distribution of DMPs across a variety of functional genomic regions is maintained across successive thresholds (Table S4).

The gene ontology (GO) and KEGG pathway enrichment for these DMPs were determined using the missMethyl package in R (26,32,37,38) which takes into account the differing number of probes per gene present on the array. Significantly enriched (FDR < 0.05) GO biological processes were subsequently summarized using REViGO which removed redundant GO terms (retained only 50% of the full list of significant terms) and visualized the remaining terms in a semantic similarity-based scatterplot (39). Semantic similarity was calculated using the simRel score, which is a functional similarity measure that ranges from 0 for terms that have no similarity to 1 for terms with maximum similarity (40).

Lastly, baboon OA-related DMPs were compared to previous findings of differential methylation associations from several human OA studies (2,4,41–49). For these comparisons, two different subsets of data were considered – (1) total data available and (2) data from matched joints and tissue types across humans and baboons (note: only knee cartilage data fit these criteria). For both data subsets, human data were limited to findings from methylation comparisons across different grades of OA (e.g., healthy vs. OA, mild OA vs. severe OA), as well as sites and genes that were also tested in the current baboon study. As the 450K array was the most common method used in human OA studies, baboon data were limited to findings from EPIC array probes that matched probes available on the 450K array. Comparisons between human and baboon findings were then done at two different levels – (1) gene-level summations of methylation patterns and (2) CpG site-specific methylation levels. For each level of comparison, the number of loci showing convergent or divergent OA methylation were identified. Loci were excluded from the comparison if conflicting methylation patterns were present across different CpG sites within a gene, across different human OA studies, across different joints, and/or across different tissue types.

In addition to DMPs, differentially methylated regions (DMRs) were also identified between each comparative group using the DMRcate package in R (50–52). This method is only concerned with the spatial proximity of loci examined and is not biased by any annotations associated with these loci. For these analyses, the individual DMP t-statistics, which were derived by fitting the M value array data to a GLM design matrix (Equations 3A) by generalized least squares using the limma package in R (10,26,27), were smoothed across each chromosome using a recommended Gaussian kernel bandwidth of 1000 base pairs with a scaling factor of 2. An expected value of this smoothed estimate with no experimental effects was also modelled using a Satterthwaite approximation (53) in order to calculate a subsequent significance test for each DMP. A default threshold was then applied to p-values adjusted for multiple testing (32) to identify FDR-corrected significant DMPs. Finally, these significant DMPs were agglomerated together into DMRs based on chromosomal location and such that each DMR contained at least 2 CpG sites that were less than 1000 base pairs apart.

#### Supplemental Figures

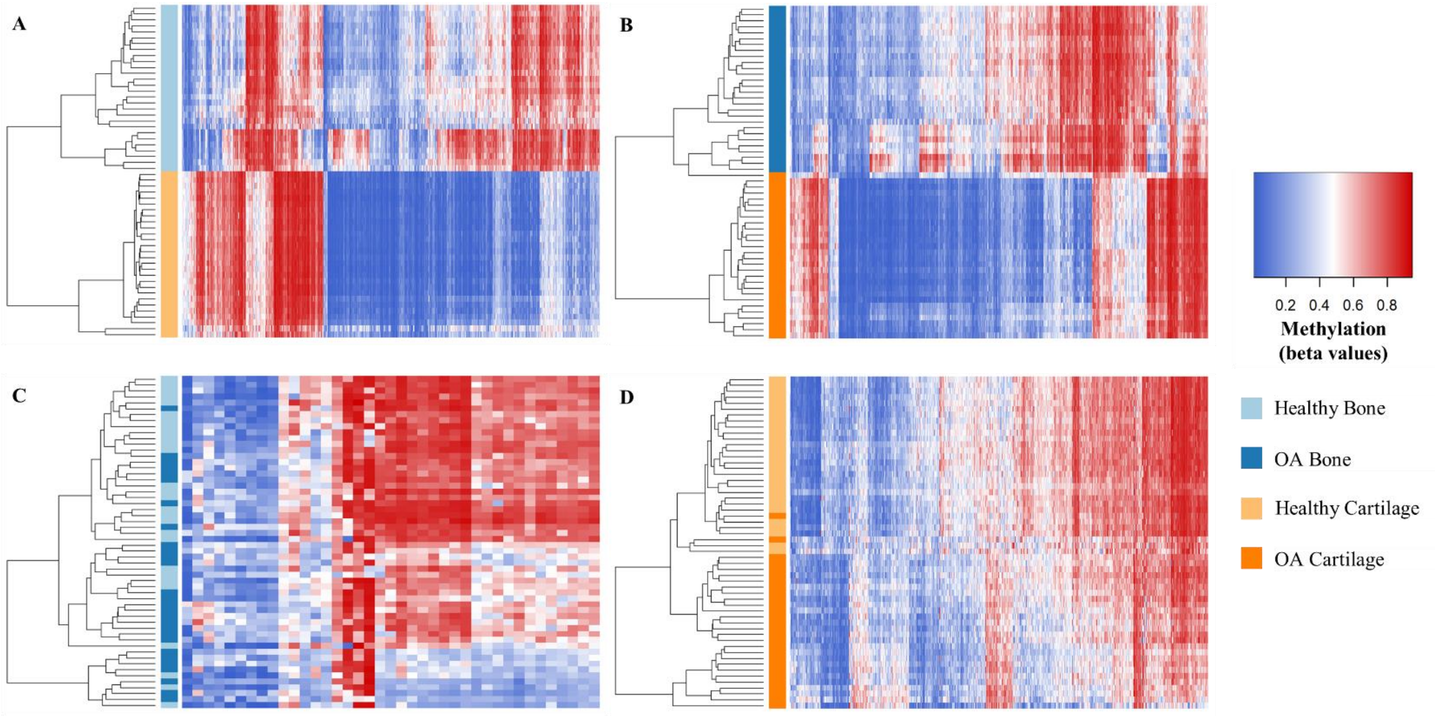

**Figure S1. Methylation Levels at DMPs Identified for Tissue Type and Disease State Comparisons.** Heatmaps depicting the DNA methylation levels (β values) of DMPs that remained significant after accounting for kinship and had at least a 10% change in mean methylation ($\Delta\beta \geq 0.1$), specifically (A) the top 15,000 DMPs between bone and cartilage (x-axis) in healthy baboons (y-axis, n=56), (B) the top 15,000 DMPs between bone and cartilage (x-axis) in OA baboons (y-axis, n=56), (C) all DMPs between OA and healthy baboons (x-axis) in bone tissues (y-axis, n=56), and (D) all DMPs between OA and healthy baboons (x-axis) in cartilage tissues (y-axis, n=56). Red indicates higher methylation at a DMP, while blue indicates lower methylation at a DMP. The dendrogram of all samples (y-axis) clusters individuals based on the similarity of their methylation patterns.

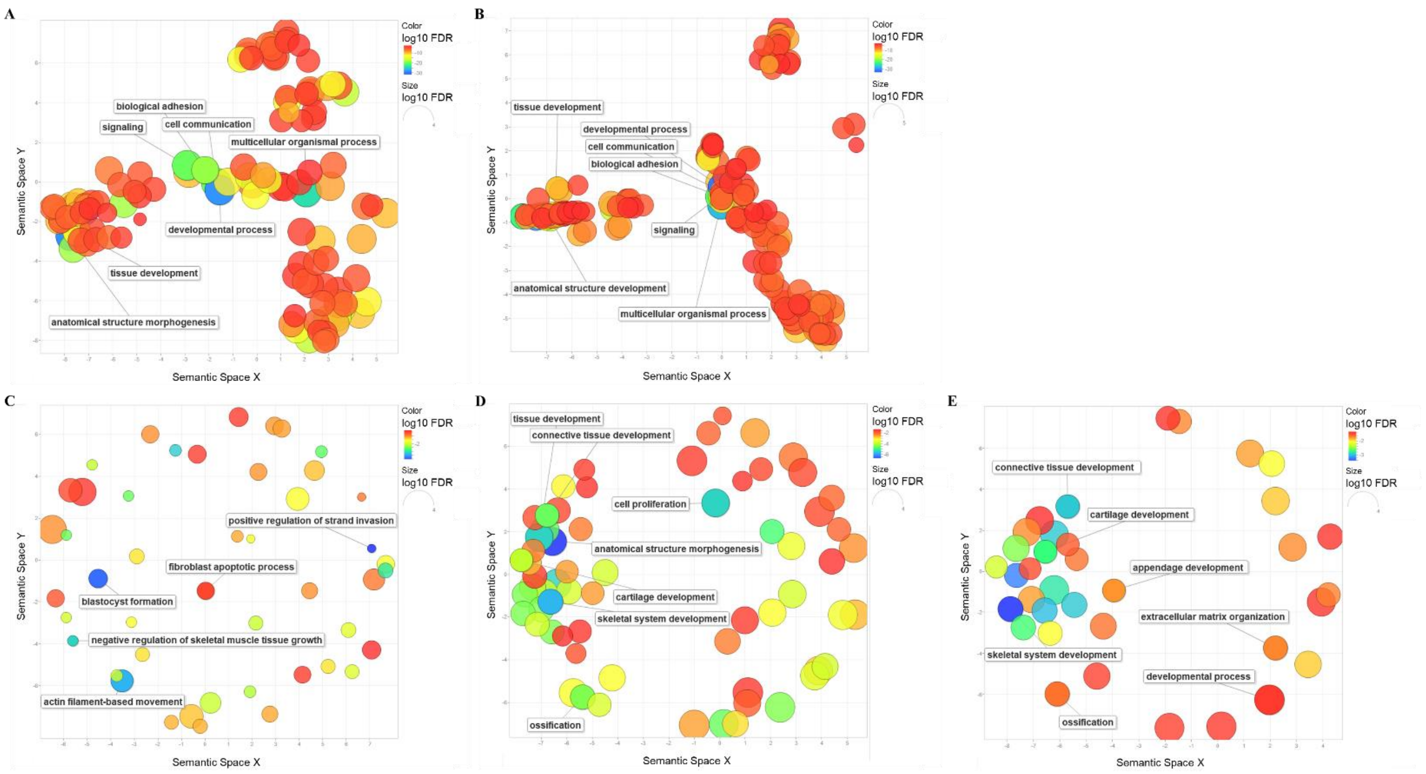

**Figure S2. Functional Enrichment of DMPs Identified for Tissue Type and Disease State Comparisons.** Multidimensional scaling plot summarizing the GO biological process terms that are significantly enriched (FDR < 0.05) for DMPs that remained statistically significant after accounting for kinship with at least a 10% change in mean methylation ($\Delta\beta \geq 0.1$) between tissue type and disease state comparisons, taking into account the differing number of probes per gene present on the EPIC array. DMPs were identified between (A) healthy bone vs. healthy cartilage, (B) OA bone vs. OA cartilage, (C) OA bone vs. healthy bone, (D) OA cartilage, and (E) all four combinations of disease state and tissue type. REViGO was used to remove redundant GO terms (retained only 50% of the full list of significant terms, see FileS3) and to visualize the remaining terms in a semantic similarity-based scatterplot. Semantic similarity was calculated using the simRel score, which is a functional similarity measure that ranges from 0 for terms that have no similarity to 1 for terms with maximum similarity. These pairwise semantic similarity scores are plotted in multidimensional scaling space such that similar GO terms are located close to one another in the plot. The color and size of each GO term are based on the log10 FDR value, and some GO terms of interest have their descriptions provided in the plot.

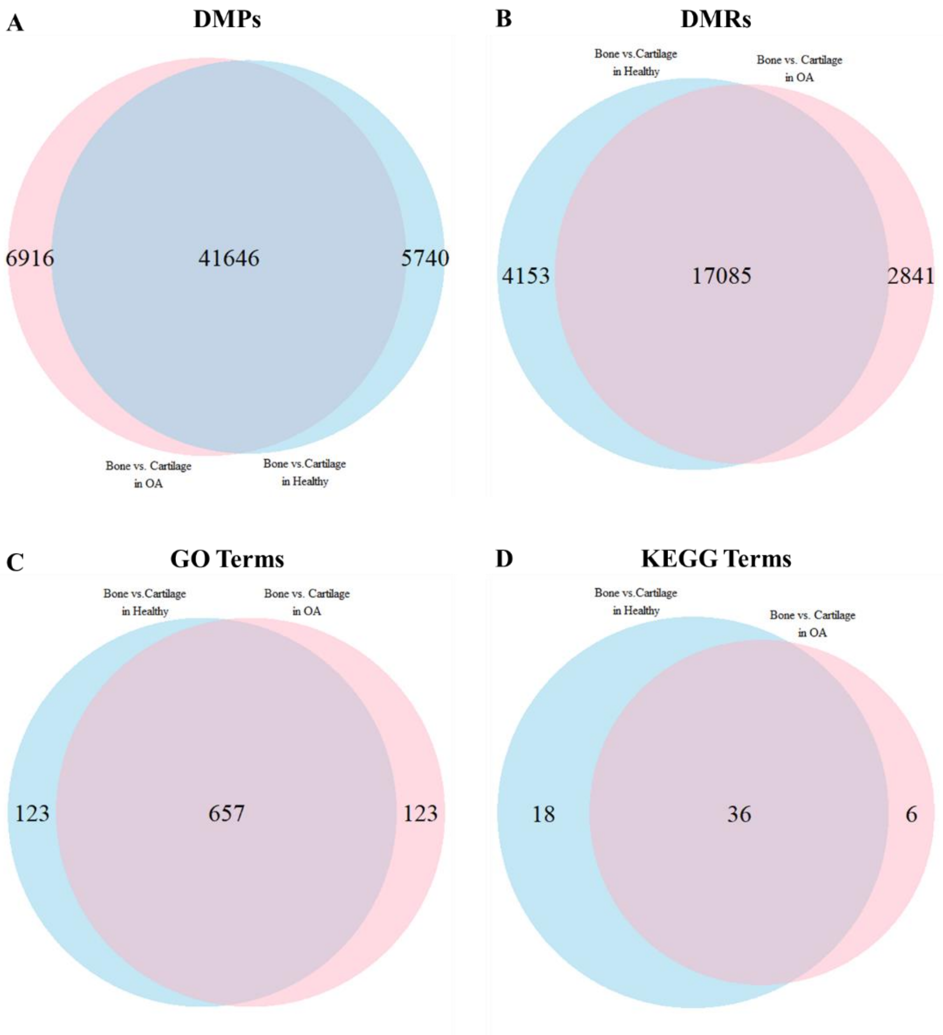

**Figure S3. Overlap of Tissue Type Differential Methylation Findings in Healthy and OA Baboons.** Venn diagrams showing (A) the overlap of DMPs that remained statistically significant after accounting for kinship with at least a 10% change in mean methylation ($\Delta\beta \geq 0.1$) between healthy bone vs. healthy cartilage and OA bone vs. OA cartilage, (B) the overlap in significant DMRs between healthy bone vs. healthy cartilage and OA bone vs. OA cartilage, (C) the overlap in GO biological process terms that are significantly enriched for DMPs that remained statistically significant after accounting for kinship with $\Delta\beta \geq 0.1$ between healthy bone vs. healthy cartilage and OA bone vs. OA cartilage, and (D) the overlap in KEGG pathways that are significantly enriched for DMPs that remained statistically significant after accounting for kinship with $\Delta\beta \geq 0.1$ between healthy bone vs. healthy cartilage and OA bone vs. OA cartilage.

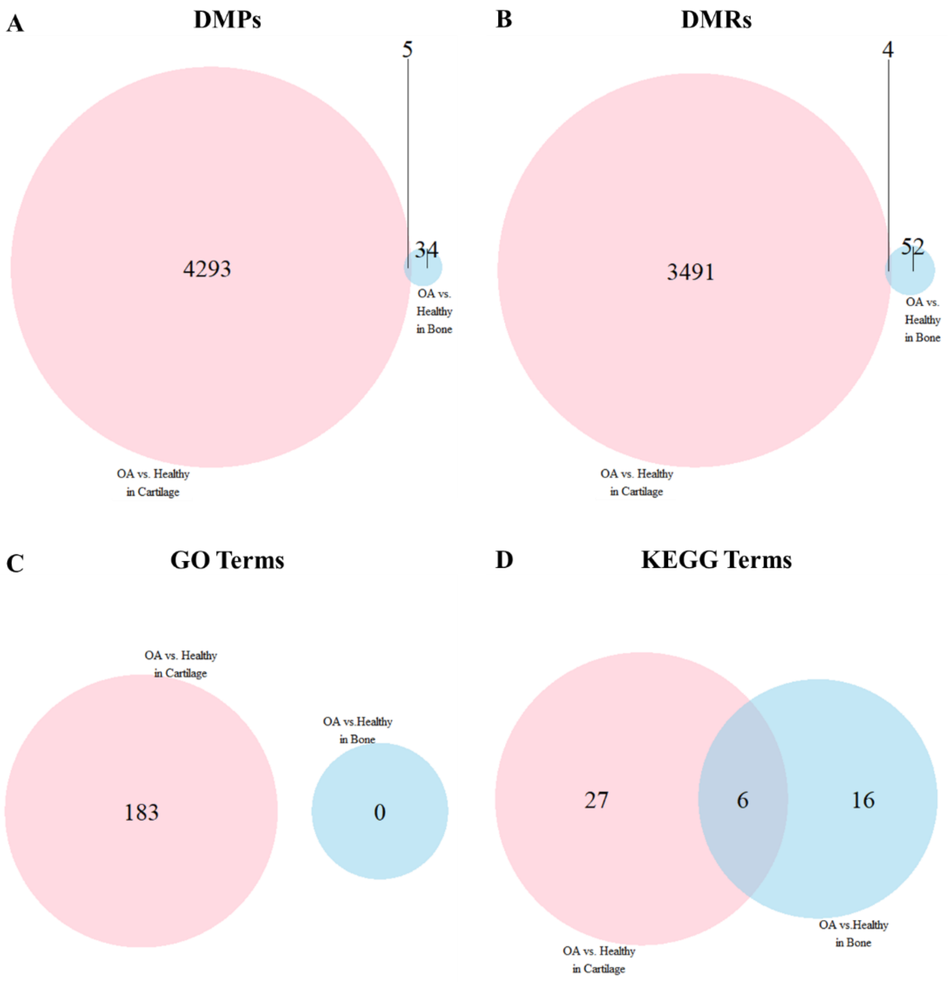

**Figure S4. Overlap of Disease Status Differential Methylation Findings in Bone versus Cartilage.** Venn diagrams showing (A) the overlap of DMPs that remained statistically significant after accounting for kinship with at least a 10% change in mean methylation ($\Delta\beta \geq 0.1$) between OA bone vs. healthy bone and OA cartilage vs. healthy cartilage, (B) the overlap in significant DMRs between OA bone vs. healthy bone and OA cartilage vs. healthy cartilage, (C) the overlap in GO biological process terms that are significantly enriched for DMPs that remained statistically significant after accounting for kinship with $\Delta\beta \geq 0.1$ between OA bone vs. healthy bone and OA cartilage vs. healthy cartilage, and (D) the overlap in KEGG pathways that are enriched for DMPs that remained statistically significant after accounting for kinship with $\Delta\beta \geq 0.1$ between OA bone vs. healthy bone and OA cartilage vs. healthy cartilage.

#### Supplemental Table Legends

All supplemental tables are provided in TableS1-S8.xlsx.

**Table S1. Baboon Sample Set.**

Details of baboon sample set, outlining each sample's animal identification number (Animal ID), species (Species), tissue type (Tissue Type), classification as healthy or having severe knee OA (Disease Status), sex (Sex), age in years (Age), and adult steady state weight in kilograms (Weight), as well as the identification number of the beadchip (Array ID) and array (Position) that each sample was run on.

**Table S2. Number of Significant DMPs Identified.**

The number of DMPs identified between comparative groups. Results include the number of DMPs that were statistically significant before accounting for kinship, the number of DMPs that remained statistically significant after accounting for kinship, and the number of DMPs that remained statistically significant after accounting for kinship with at least a 10% change in mean methylation ($\Delta\beta \geq 0.1$) between comparative groups. Before accounting for kinship, 66.4% (127,519) of probes were differentially methylated between bone and cartilage in healthy baboons, 62.6% (120,064) between bone and cartilage in OA baboons, 0.2% (384) between OA and healthy bone, 11.1% (21,372) between OA and healthy cartilage, and 2.7% (5,136) between all four combinations of disease state and tissue type. After accounting for genetic relatedness, 2.2% (2,807), 2.0% (2,419), 0.5% (2), 0.8% (172), and 0.5% (24) of the originally identified DMPs do not maintain significant methylation associations, respectively. Of those DMPs that remained statistically significant after accounting for kinship, only 38.0% (47,386), 41.3% (48,562), 10.2% (39), 20.3% (4,298), and 52.4% (2,676) had $\Delta\beta \geq 0.1$, suggesting that these DMPs may have regulatory and biological effects. See FileS1 for more details.

**Table S3. Direction of Change in Significant DMPs.**

The number of DMPs that show significant differential hypo-methylation (negative) or hyper-methylation (positive) between comparative groups. Results include the number of DMPs that were statistically significant before accounting for kinship, the number of DMPs that remained statistically significant after accounting for kinship, and the number of DMPs that remained statistically significant after accounting for kinship with at least a 10% change in mean methylation ($\Delta\beta \geq 0.1$) between comparative groups. See FileS1 for more details.

**Table S4. Genomic Distribution of Significant DMPs.**

Details on the number of DMPs that occupy different genomic regions (based on the human genome hg19). Results include the number of DMPs that were statistically significant before accounting for kinship, the number of DMPs that remained statistically significant after accounting for kinship, and the number of DMPs that remained statistically significant after accounting for kinship with at least a 10% change in mean methylation ($\Delta\beta \geq 0.1$) between comparative groups. Number of Genes indicates the number of unique gene symbols associated with DMPs. Probes Per Gene indicates the average number of DMPs within each associated gene. For genomic locations, TSS200 and TSS1500 indicate the transcription start site areas between the start of the gene and 200bp upstream or 1500bp upstream respectively, 5'UTR and 3'UTR indicate the untranslated regions of genes, 1st Exon indicates the first exon within a gene, ExonBnd indicates the gene body exons, and Gene Body indicates any area within the exons and introns of a gene. For proximity to CpG islands, Island indicates areas within CpG islands, North Shelf indicates areas 2-4kb upstream from a CpG island, North Shore indicates areas up to 2kb upstream from a CpG island, South Shelf indicates areas 2-4kb downstream from a CpG island, South Shore indicates areas up to 2kb downstream from a CpG island, and Open Sea indicates isolated CpG site in the genome.

**Table S5. Number of Significant DMRs Identified.**

The number of significant DMRs identified between comparative groups, along with the number of unique gene names that overlapped with these DMRs, the average, minimum, and maximum number of CpGs per DMR, the average, minimum, and maximum length of DMRs, and gene symbol names associated with the top 10 DMRs for each comparative group. See FileS2 for additional details.

**Table S6. Comparison of OA-Related Methylation between Humans and Baboons.**
The number of OA-related differentially methylated loci that overlap in humans and baboons. Comparisons between human and baboon findings were done at two different levels (Level of Comparison) – (1) gene-level summations of methylation patterns and (2) CpG site-specific methylation levels. Two different subsets of data were considered for comparisons (Data Compared) – (1) total data available and (2) data from matching joints and tissue types across humans and baboons (note: only knee cartilage data fit these criteria). For both data subsets, human data only included findings from methylation comparisons across different grades of OA (e.g., healthy vs. OA, mild OA vs. severe OA) that were also tested in the current baboon study, and baboon data only included findings from EPIC array probes that matched probes available on the 450K array, which was the most common method used in human OA studies. For each level of comparison, the number of loci showing the same patterns of OA methylation between humans and baboons (Conserved) or opposite patterns of OA methylation between humans and baboons (Divergent) are listed. Additionally, the number of OA-related loci excluded from the comparison due to conflicting methylation patterns across different CpG sites within a gene, across different human OA studies, across different joints, and/or across different tissue types (Excluded) are provided. Further, the number of OA-related loci identified as significant in humans but non-significant in baboons (Lacking Baboon DMP) and the number of OA-related loci identified as significant in baboons but non-significant in humans (Lacking Human DMP) are indicated. See Table S7 and FileS5 for additional details.

**Table S7. Details of Gene-Level and CpG-Level Comparisons between Humans and Baboons.**
Details of the OA-related differentially methylated loci that overlap in humans and baboons. Comparisons between human and baboon findings were performed in four different subsets of data – (1) gene-level summations of methylation patterns from all available data merged across different joints (e.g., knee, hip) and tissue types (e.g., bone, cartilage) (Table S7a), (2) gene-level summations of methylation patterns from joint- and tissue-matched data across humans and baboons (note: only knee cartilage data fit these criteria) (Table S7b), (3) CpG site-specific methylation levels from all available data merged across different joints and tissue types (Table S7c), and (4) CpG site-specific methylation levels from joint- and tissue-matched data across humans and baboons (Table S7d). For each comparison, the genes (Gene) and/or the CpG sites (CpG) with OA-related changes in methylation are listed. When relevant, the skeletal element (Joint) and tissue type (Tissue) from which data were collected are also noted. Additionally, the pattern of methylation comparatively observed in OA is provided for each species (OA Methylation), as well as how the patterns of methylation overlap between species (Evolutionary Comparison) – similar patterns of OA methylation between species (Conserved), opposite patterns of OA methylation between species (Divergent), patterns of OA methylation that are conflicting and do not show a uniform signal (Excluded), patterns of OA methylation significant in humans but non-significant in baboons (NoBaboonDMP), or patterns of OA methylation significant in baboons but non-significant in humans (NoHumanDMP). See FileS5 for additional details.

**Table S8. Comparative OA-Related Differential Methylation Results for Example Genes.**
Comparative (baboon vs. human) OA-related differential methylation results for three genes – *TBX4*, *HOXD8*, and *RUNX1*. Table includes gene-level and CpG-site-level data used in baboon and human comparisons. Specifically, each candidate gene is listed (Gene) along with the identification number of each CpG site examined (Array Probe ID) and whether the gene/CpG was tested in baboons (Tested in Current Baboon Study?) or in previous human OA studies (Tested on 450K?). Annotation information for each CpG site (Human Gene Symbol, Human Chromosome, Human CpG Position, Baboon Gene Symbol, Baboon Chromosome, Baboon CpG Position) and the average β values for each baboon comparative group (Average β Values) are also provided. Loci found to be significantly differentially hypo- or hyper-methylated in OA cartilage or OA bone as compared to healthy tissue are noted for baboons (Significant OA-Related Methylation Change in Baboons) and for humans (Significant OA-Related Methylation Change in Humans). In baboons, all DMPs listed are DMPs that remained statistically significant after accounting for kinship, and DMPs with at least a 10% change in mean methylation ($\Delta\beta \geq 0.1$) between comparative groups are shown in bold. None of the listed probes that were tested in baboons contain potential baboon SNP variants at the CpG site. In humans, the reference for each OA-related methylation change is noted. See FileS5 for additional details.

#### Supplemental File Descriptions

All supplemental files are provided in FileS1-S5.zip.

**FileS1_DMP.xlsx**

Gene details of the significant DMPs identified. Tables list the significant DMPs (Adjusted P-Value < 0.05) between each comparative group – healthy bone vs. healthy cartilage (Bone vs Cartilage in Healthy), OA bone vs. OA cartilage (Bone vs Cartilage in OA), OA bone vs. healthy bone (OA vs Healthy in Bone), OA cartilage vs. healthy cartilage (OA vs Healthy in Cartilage), and all four combinations of disease state and tissue type (All Four Combinations). Tables include the identification number of each DMP probe (EPIC Array Probe ID), as well as additional annotation information (Human Gene Symbol, Baboon Gene Symbol, Baboon Chromosome, Baboon CpG Position), the average β values for each comparative group (Average β Values), and the difference in β values across groups (Absolute Difference in β Values). Results for the initial DMP analysis before accounting for kinship (DMP Analysis) includes the log fold difference in M values between each comparative group (Log Fold Change in M Values) and the p-values for each DMP after accounting for multiple testing (Adjusted P-Value). Results for the DMP analyses that account for kinship (Kinship Analysis) include the log likelihood values when the comparative group variables of interest were included in the GLMM (Log Likelihood with Variable of Interest), the log likelihood values when the comparative group variables of interest were excluded from the GLMM (Log Likelihood without Variable of Interest), the chi-square value (X2), and the p-value for these tests (P-Value). The table order is based on the kinship analysis p-values (smallest to largest) and divided so that DMPs with at least a 10% change in mean methylation ($\Delta\beta \geq 0.1$) between comparative groups and DMPs with $\Delta\beta < 0.1$ are separated.
*Probes that contain potential baboon SNP variants at the CpG site (722 probes in DMPs between bone and cartilage in healthy baboons, 693 probes in DMPs between bone and cartilage in OA baboons, 2 probes in DMPs between OA and healthy bone, 115 probes in DMPs between OA and healthy cartilage, and 25 probes in DMPs between all four combinations of tissue type and disease state).

**FileS2_DMR.xlsx**

Gene details of the significant DMRs identified. Tables list the significant DMRs (Stouffer Transformed FDR < 0.05) between each comparative group – healthy bone vs. healthy cartilage (Bone vs Cartilage in Healthy), OA bone vs. OA cartilage (Bone vs Cartilage in OA), OA bone vs. healthy bone (OA vs Healthy in Bone), OA cartilage vs. healthy cartilage (OA vs Healthy in Cartilage), and all four combinations of disease state and tissue type (All Four Combinations). These analyses used a Gaussian kernel bandwidth of 1000 base pairs with a scaling factor of 2 as recommending in the DMRcate package in R. DMRs had to contain at least 2 CpG sites that were less than 1000 base pairs apart, and p-values were adjusted using the Benjamini-Hochberg method. Tables include the genomic location of each DMR based on the human genome (Human Chromosome, Human DMR Start, Human DMR End), the length of each DMR in base pairs (DMR Length (bp)), the number of CpG sites constituting the significant region (No. CpGs), the minimum adjusted p-values from the CpGs constituting each DMR (Adjusted P-Value), the stouffer transformations of the groups of false detection rates for individual CpG sites as DMR constituents for each DMR (Stouffer Transformed FDR), the maximum absolute beta fold change within each DMR (Max. Log Fold Change in β Values), the mean beta fold change within each DMR (Mean Log Fold Change in β Values), and lists of genes with promotor regions that overlap with each DMR (Overlapping Promotors).

**FileS3_GO.xlsx**

GO biological processes enriched for significant DMPs. Tables contain the GO biological process terms that are significantly enriched (FDR < 0.05) for DMPs that remained statistically significant after accounting for kinship with at least a 10% change in mean methylation ($\Delta\beta \geq 0.1$) between comparative groups – healthy bone vs. healthy cartilage (Bone vs Cartilage in Healthy), OA bone vs. OA cartilage (Bone vs Cartilage in OA), OA bone vs. healthy bone (OA vs Healthy in Bone), OA cartilage vs. healthy cartilage (OA vs Healthy in Cartilage), and all four combinations of disease state and tissue type (All Four Combinations) – taking into account the differing number of probes per gene present on the EPIC array. Tables include the identification numbers (GO ID) and terms (GO Biological Process Term) for each significantly enriched GO term, the total number of genes associated with each GO term (No. Genes Total), the number of genes with significant DMPs that are also associated with each GO term (No. Gene with DMPs), the p-value for over-representation of each GO term (P-Value), and the false discovery rate for each GO term (FDR). For OA bone vs. healthy bone (OA vs Healthy in Bone), no GO categories were significant at 5% FDR, so all GO functions with p-values < 0.05 are listed.

**FileS4_KEGG.xlsx**
KEGG pathways enriched for significant DMPs. Tables contain the KEGG pathways that are significantly enriched (FDR < 0.05) for DMPs that remained statistically significant after accounting for kinship with at least a 10% change in mean methylation ($\Delta\beta \geq 0.1$) between comparative groups – healthy bone vs. healthy cartilage (Bone vs Cartilage in Healthy), OA bone vs. OA cartilage (Bone vs Cartilage in OA), OA bone vs. healthy bone (OA vs Healthy in Bone), OA cartilage vs. healthy cartilage (OA vs Healthy in Cartilage), and all four combinations of disease state and tissue type (All Four Combinations) – taking into account the differing number of probes per gene present on the EPIC array. Tables include the identification numbers (KEGG ID) and pathways (KEGG Pathway) for each significantly enriched KEGG term, the total number of genes associated with each KEGG pathway (No. Genes Total), the number of genes with significant DMPs that are also associated with each KEGG pathway (No. Gene with DMPs), the p-value for over-representation of each KEGG pathway (P-Value), and the false discovery rate for each KEGG pathway (FDR). For OA bone vs. healthy bone (OA vs Healthy in Bone), OA cartilage vs. healthy cartilage (OA vs Healthy in Cartilage), and all four combinations of disease state and tissue type (All Four Combinations), no KEGG pathways were significant at 5% FDR, so all KEGG pathways with p-values < 0.05 are listed.

**FileS5_Compare.xlsx**
Comparison of OA-related differential DNA methylation in humans and baboons. The HumanOA and BaboonOA tables outline details for several genes and CpG sites that have been previously identified as differentially methylated in human OA studies and the current baboon OA study, respectively. The References Table lists the references from which previous human OA methylation findings were compiled.

In the HumanOA table, genes with OA-related changes in methylation (Gene) and, when available, information regarding the locus identification number that was differentially methylated (Array Probe ID) are listed. Regarding human OA findings, the publication source of each finding (Reference), the method used to collect data (Method), the specific OA phenotype evaluated (Phenotype of Interest), the general phenotypic comparison examined (Comparison), and the skeletal element (Joint) and tissue type (Tissue) collected are provided. When available, the average methylation level of the OA phenotype (β Value for OA) is also noted. Additionally, the pattern of methylation comparatively observed in the OA phenotype (OA Methylation Level) is provided, along with details on known functional characterizations of loci (Functional Characterization) from different literature sources (Functional Reference). Information on whether a CpG site was also tested in the current baboon study (out of the 191,954 finalized probe set) (CpG Tested in Current Baboon Study?) or whether a gene was also tested in the current baboon study (Gene Tested in Current Baboon Study?) is also noted, as well as corresponding details from the current baboon study – EPIC array probe annotation information (Baboon Gene Symbol, Baboon Chromosome, Baboon CpG Position) and the average β values for each baboon disease and tissue comparative group (OA Bone, Healthy Bone, OA Cartilage, Healthy Cartilage).

In the BaboonOA table, genes with OA-related changes in methylation (Gene) and information regarding the locus identification number that was differentially methylated (Array Probe ID) are listed. For each locus, the type of DMP detected is specified – DMP that was statistically significant before accounting for kinship (DMP), DMP that remained statistically significant after accounting for kinship (Kinship DMP), and/or DMP with at least a 10% change in mean methylation between comparative groups ($\Delta\beta \geq 0.1$ DMP). A summary of these DMP types is also provided (DMP Summary). Additionally, the phenotypic comparison examined (Comparison), the skeletal element (Joint) and tissue type (Tissue) collected, the pattern of methylation comparatively observed in the OA phenotype (OA Methylation Level), and the significance level of the association (Adjusted P-Value) are listed. Information on whether EPIC array probes match those available on the 450K array is also noted (CpG Tested on 450K?), as this was the primary method used in human OA studies. Corresponding details from the current baboon study – probe annotation information (Baboon Gene Symbol, Baboon Chromosome, Baboon CpG Position) and the average β values for each baboon disease and tissue comparative group (OA Bone, Healthy Bone, OA Cartilage, Healthy Cartilage) – are also provided.

Abbreviations: NA = gene symbol not available, 27K = Illumina Infinium HumanMethylation27K BeadChip, 450K = Illumina Infinium HumanMethylation450K BeadChip, microarray = Agilent Human Promoter Microarray, OA = osteoarthritis, OP = osteoporosis.
